# A Statistical Model of Shared Variability in the Songbird Auditory System

**DOI:** 10.1101/113670

**Authors:** Lars Buesing, Ana Calabrese, John P. Cunningham, Sarah M. N. Woolley, Liam Paninski

## Abstract

Vocal communication evokes robust responses in primary auditory cortex (A1) of songbirds, and single neurons from superficial and deep regions of A1 have been shown to respond selectively to songs over complex, synthetic sounds. However, little is known about how this song selectivity arises and manifests itself on the level of networks of neurons in songbird A1. Here, we examined the network-level coding of song and synthetic sounds in A1 by simultaneously recording the responses of multiple neurons in unanesthetized zebra finches. We developed a latent factor model of the joint simultaneous activity of these neural populations, and found that the shared variability in the activity has a surprisingly simple structure; it is dominated by an unobserved latent source with one degree-of-freedom. This simple model captures the structure of the correlated activity in these populations in both spontaneous and stimulus-driven conditions, and given both song and synthetic stimuli. The inferred latent variability is strongly suppressed under stimulation, consistent with similar observations in a range of mammalian cortical regions.

## 1 Introduction

Vocal communicators depend on auditory perception for survival and reproduction. Songbirds rely on auditory processing to learn song, recognize individuals, and judge the fitness of potential mates (Catchpole and Slater, 2004). Electrophysiological studies on the firing properties of neurons (recorded in brain regions that involve song perception) have typically recorded from one neuron at a time (Theunissen et al., 2004, Margoliash, 1997, Doupe and Konishi, 1991). Using this approach, auditory neurons that respond more strongly to vocalizations than to similarly complex, non-natural sounds have been found (Leppelsack, 1978, Leppelsack and Vogt, 1976, Leppelsack, 1983, Grace et al., 2003), and changes in single-neuron representations of songs versus other complex sounds are well documented (Woolley et al. 2005, Schneider and Woolley, 2011, Jeanne et al., 2011, Meliza and Margoliash, 2012).

While a great deal has been discovered with single-neuron recordings, these studies cannot elucidate the structure of correlated dynamics in neuronal networks. Recent studies have demonstrated the importance of neuronal network properties in gaining a better understanding of how the brain transforms sensory signals into information-bearing percepts (Schneidman et al., 2006, Jones et al., 2007, Pillow et al., 2008, Yu et al., 2009, Paninski et al., 2010, Berkes et al., 2011, Macke et al., 2011, Buesing et al., 2014, Cunningham and Yu, 2014, Okun et al., 2015). These studies make use of the fact that simultaneously recorded cells generically show dependent trial-to-trial and/or temporal activity fluctuations, called *shared variability* (Sakata and Harris, 2009, Churchland et al., 2010, 2012, Smith et al., 2013, Hansen et al., 2012). Shared variability is shaped by a large number of phenomena, such as common, unobserved inputs originating from presynaptic neurons (Zohary et al., 1994, Shadlen and Newsome, 1998) or by modulatory states generated internally, like attention, arousal, or adaptation (Cohen and Newsome, 2008, Nienborg and Cumming, 2009, Ecker et al., 2010, 2014,Harris and Thiele, 2011). Shared variability has been shown in some cases to reflect the anatomical connectivity of the underlying circuitry (Gerhard et al., 2013, Okun et al., 2015). A detailed characterization of shared variability is crucial for understanding of neural coding as it can dramatically influence the fidelity of a population code (Averbeck et al., 2006Cohen and Kohn, 2011). Analyzing and modeling shared variability therefore holds promise for uncovering potential mechanisms of stimulus selectivity in sensory areas such as the avian A1.

When analyzing multi-dimensional activity recordings, we would ideally characterize and quantify shared variability and decompose it into separate sources, corresponding to distinct physiological mechanisms, which are superimposed in the raw data. These desiderata cannot be met by simple analyses exclusively based on noise correlations, highlighting the need for improved data analysis techniques. Factor models, which are generalizations of Principal Component Analysis and classical Factor Analysis (PCA and FA; see (Cunningham and Ghahramani, 2015) for a review), have proven very useful in this context, allowing one to identify anatomically defined cell types (Buesing et al., 2014), to identify task-related processing stages of network activity (Petreska et al., 2011), to characterize the effect of anesthesia or attention on population firing (Ecker et al., 2014, Rabinowitz et al., 2015), and to show that the relative contribution of different sources of variability can vary across brain regions (Goris et al., 2014).

Here we study the multi-dimensional structure of neural activity in the zebra finch primary auditory cortex observed with multi-electrode extra-cellular recordings. By applying a latent factor model, we perform a detailed analysis of the shared variability and show that it has (to good approximation) only one degree-of-freedom, suggesting that trial-to-trial variability in the recordings can be interpreted as generated by an unobserved network state that influences simultaneously recorded neurons in a highly coordinated way.

## 2 Materials and Methods

### 2.1 Stimuli

Two classes of sound stimuli were used: the songs of 20 adult male zebra finches and 10 unique samples of modulation-limited (ML) noise. ML noise is correlated Gaussian noise designed to match song stimuli in power, frequency range (250-8000 Hz), and maximum spectral and temporal modulation frequencies (Schneider and Woolley, 2011). Each ML noise stimulus was 2 seconds in duration. Stimuli were delivered free-field through a flat frequency response speaker positioned 20 cm in front of the bird, at a mean intensity of 65 dB sound pressure level (SPL). Between 30 and 40 response trials were obtained for each of the 10 ML noise stimuli. Between 15 and 20 trials were obtained for each of the 20 songs. Trials for different stimuli were presented in pseudo-random order. Inter-trial intervals were determined by randomly sampling from a uniform distribution between 2 s and 3 s.

### 2.2 Electrophysiology

Two days before recordings, birds were anesthetized with a single intramuscular injection of 0.04 ml of Equithesin and placed in a custom-designed stereotaxic holder. Craniotomies were made 1.3 mm lateral and 1.3mm anterior from the bifurcation of the midsagittal sinus (stereotactic coordinates). For each bird, a small metal post was then affixed to the skull using dental acrylic, and a grounding wire was cemented in place with its end just beneath the skull, approximately 5 – 10 mm lateral to the junction of the midsagittal sinus. Birds recovered from surgery for two days. Recordings were made in A1 regions CLM and Field L subregions L1, L, L2a, L2b, and L3 in head-fixed, unanesthetized, male zebra finches (*Taeniopygia guttata, n* = 6). Recordings were made using a planar multichannel silicon polytrode (4×4 electrode layout, 177 *μ*m^2^ contact surface area, 100 *μ*m inter-contact distance in the dorsal-ventral direction and 125*μ*m inter-contact distance in the anterior-posterior direction; NeuroNexus Technologies). Each region was recorded in 4, 5 or 6 out of 6 birds (CLM: birds 1, 3, 4, and 5; L1: birds 1, 2, 4, and 5; L2a: birds 1 to 5; L2b: birds 1 to 6; L: birds 2 to 6; L3: birds 1 to 5), yielding 219 single units recorded from superficial regions CLM and L1 (CLM: 153, L1: 66), 377 from intermediate regions L2a, L2b and L (L2a: 47, L2b: 47, L: 283), and 237 from the deep region L3. For each animal, recordings were performed daily over approximately one week, and each recording session was approximately 6 hrs long. Signals were amplified and band-pass filtered between 300 and 5000 Hz, digitized at 25 kHz (RZ5; Tucker-Davis Technologies), and stored for off-line processing.

### 2.3 Data Analysis

#### 2.3.1 Spike sorting

Data analysis was carried out in MATLAB (Mathworks). Spikes were sorted offline with the automated sorting algorithm WaveClus (Quiroga et al., 2004). First, a nonlinear filter that increases the signal-to-noise ratio (by emphasizing voltage deflections that are both large in amplitude and high in frequency content) was applied to the bandpass filtered voltage trace of each channel (Kim and Kim, 2000, Hill et al., 2011). Second, spikes were detected and sorted in an unsupervised manner by WaveClus. Third, the output of this algorithm was refined by hand for each electrode, taking into account the waveform shape and interspike interval distribution. To quantify the quality of the recording we computed the signal-to-noise (SNR) of each candidate unit as the ratio of the average waveform amplitude to the SD of the waveform noise (Hill et al., 2011, Kelly et al., 2007). Only candidate units that had an SNR greater than 3.5 were considered single units and were included in the analyses (median SNR across all included units = 7.47). This procedure yielded a total of 833 units recorded during 96 recording sessions from 6 birds.

### 2.4 Poisson linear dynamical system model

Here we give the detailed definition of the Poisson Linear Dynamical System model (Fig. 1) and describe the associated inference and parameter estimation methods. For the sake of simplicity, we break the exposition into two parts. We first describe the *peristimulus time histogram* (PSTH) model that captures the average evoked neural activity. Second, we discuss the full Poisson linear dynamical system model (PLDS,Macke et al. (2011)) that also captures trial-to-trial variability around the average responses.

**Figure 1:**
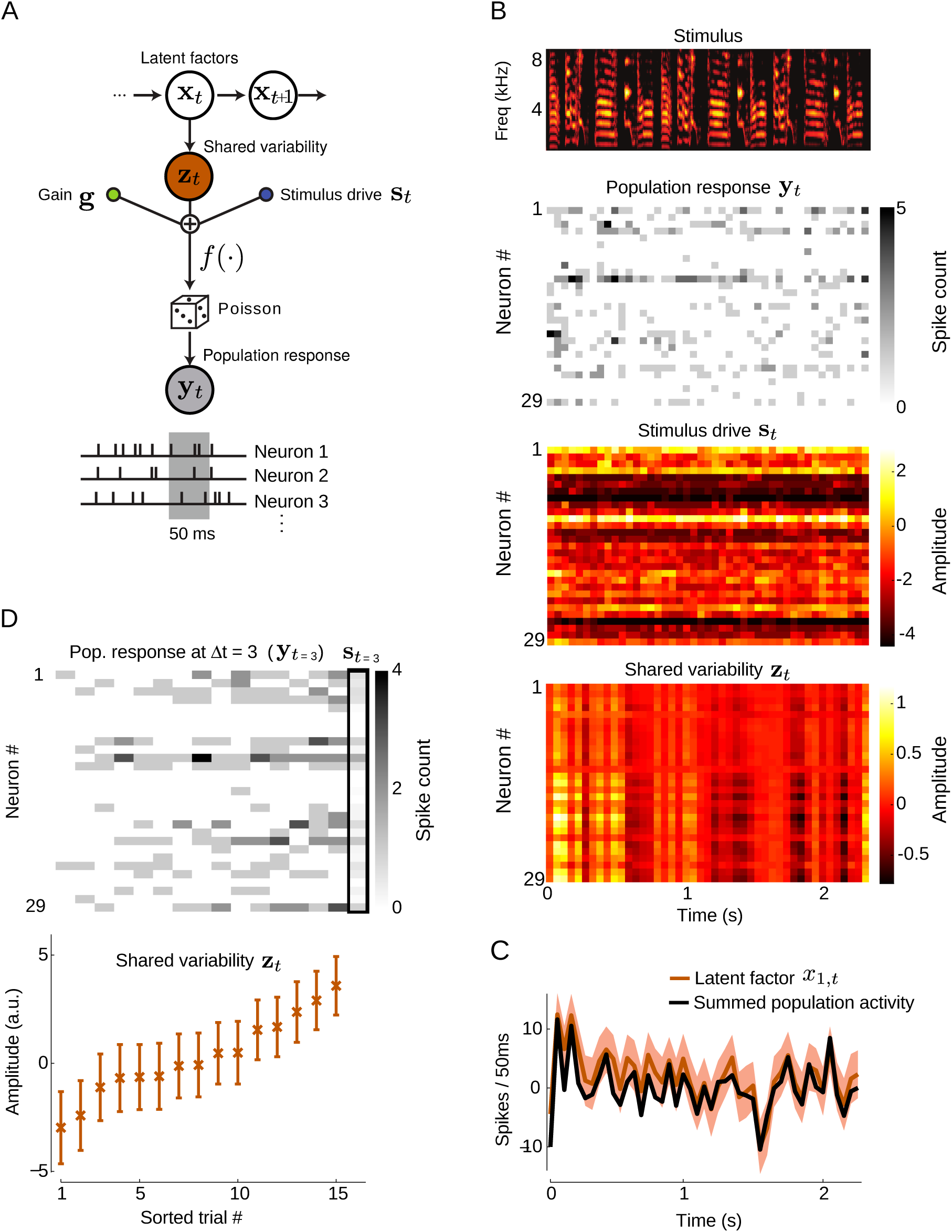
Latent factor model for analyzing shared variability. (**A**) Schematic of the PLDS model. Single-trial population spike count responses **y**_t_ (in 50 ms) are modeled as Poisson distributed with a rate that depends on three contributions: I) an external stimulus drive **s**_t_, II) a slowly varying gain-modulation g, and III) a shared variability term **z**_t_, which models fast shared variability in the neural responses caused by unobserved processes. We model the variability **z**_t_ across the population using a latent state vector **x**_t_, which compactly summarizes activity variability shared across neurons. **x**_t_ evolves in time according to linear Gaussian dynamics. (**B**) Results of applying the PLDS model to an example data set from the deep region of avian A1. (Top) Spectrogram of a zebra finch song that was used as auditory stimulus. (Second from top) Spike count responses of all simultaneously recorded cells to one presentation of the song stimulus. (Bottom) Estimated stimulus drive st and inferred posterior mean of shared variability **z**_t_. (C) Population response summed across neurons for the same trial as in (B). Shown is the deviation (black line) on the example single trial from the (time-dependent) trial average. It is highly similar to the first latent factor *x*_1,*t*_ (brown, scaled for comparison, shaded envelope is the posterior standard deviation). (D) Top: The spike count vector **y**_*t*_ at a fixed time bin *t* (*t* = 3 in this example) is shown for all 15 presentations of the song stimulus from (B), illustrating population response variability across trials. For comparison, the trial average (boxed) is shown on the right. Bottom: Inferred first factor *x*_1,*t*_ (posterior mean, errorbars are posterior standard deviation) for the corresponding spike count vectors **y**_*t*_ shown above. For both panels (top and bottom) trials were sorted with respect to the magnitude of *x*_1,*t*_.

To begin, we need to define some notation. For each of the 30 different stimuli (song and ML noise), we here considered 15 trials. Each data set also included 450 trials consisting of 500 ms of spontaneous neural activity without auditory stimulation. We binned the recorded data into spike counts in windows of size 50 ms. For a given data set, we denote with *y_kt_^mn^* the spike count of neuron *k* = 1,…, *K* (where *K* ranged from 5 to 29 across data sets) during the presentation of stimulus m = *1,…,M* (where *m* = 1,…, 20 are song, *m* = 21,…, 30 are ML noise stimuli and *m* = 31 for spontaneous activity in time bin *t* = 1,…, *T_m_* (where 52 < *T_m_* < 69 for song/ML noise stimuli and *T*_31_ = 10) on trial *n* = 1,…, *N* (with *N* =15 for song and ML noise stimuli and *N* = 450 for silence).

#### 2.4.1 PSTH model

First, we fitted a simple model to capture average evoked responses across trials as well as slowly varying aspects of the data. For each data set, we modeled the spike count *y_kt_^mn^* as a Poisson random variable with rate *f*(*λ_kt_^mn^*):

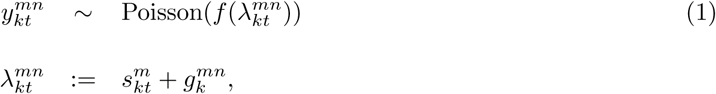

The signal drive terms *s_kt_^m^* are the same (i.e. constant) for all across trials n and capture the trial-independent influence of the stimulus on the recorded activity. The gain modulation *g_kt_^mn^* is constant during each stimulus presentation, i.e. independent of time bin *t*, and therefore captures slowly varying aspects of the data. These signal and slow gain modulation terms together constitute the *PSTH model.* The non-negative function *f* is termed the *transfer function.* In the context of generalized linear models it is generally referred to as the *inverse link function.* In the following we will use:

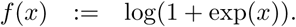

In section 2.4.11 we discuss the choice of transfer function in greater detail.

We use the notation 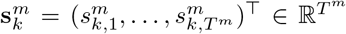 and 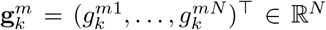. We fitted the parameters to the data using *l*_2_-regularized maximum likelihood estimation. For each neuron *k* and stimulus m we optimized the penalized log-likelihood log *p*(**y**) =: *L*_PSTH_ of the data under the model:

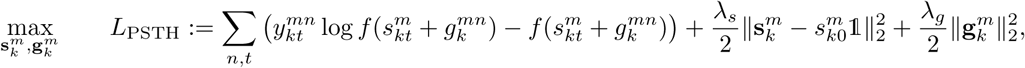

where | ∙ |_2_ is the Euclidean norm, 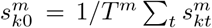 and 1 is the *T^m^*-dimensional vector with all entries equal to 1. We optimized the cost function using the l-BFGS pseudo-Newton method (Boyd and Vandenberghe, 2004). The cost function is concave, as the transfer function *f* is convex, log-concave, and monotonically increasing, guaranteeing that standard optimization methods will find the unique global maximum (Paninski, 2004).

We determined the penalty parameters *λ*_*s*_ and *λ*_*g*_ by a grid search on the five-fold cross-validated data likelihood, yielding λ_*s*_ = 1.5 and *λ*_*g*_ = 10. Given the optimal *λ*_*s*_, *λ*_*g*_ we re-fitted *s_k_^m^*, *g_k_^m^* to the complete data set. We denote the PSTH model parameters by *θ*_PSTH_ = (**s**, **g**).

#### 2.4.2 PLDS model

The PSTH model described above captures average neural evoked responses as well as slow response variability on time scales larger than a trial. To also capture neural response variability around the PSTH on shorter time scales we added an additional term *z_kt_^mn^* to the model described in eqn. (1):

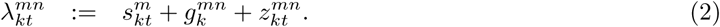

If no restrictions are placed on **z**, this model would overfit: each *z_kt_^mn^* could be chosen to explain each corresponding observed spike count *y_kt_^mn^*. As we were interested in analyzing variability that is shared across neurons, we restricted the vector 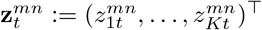 by modeling it as a linear function of a smaller number *d* of shared latent *factors* 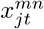 for *j* = 1,…, *d.* Using the notation 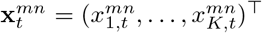, we write:

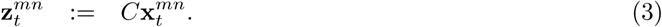

Eqn. (3) models the variability as being low-rank or low-dimensional, i.e. the variability has support on a *d*-dimensional subspace spanned by the columns of the *loading* matrix *C* ∈ ℝ ^K×d^. The vector *x_t_^mn^* was modeled as a multivariate gaussian random vector. Because each component of **x** in general influences all neurons (to the extent determined by *C*), this term imparts the shared portion of the data variability; as such we will hereafter refer to **z** as the shared variability term. From now on we will drop the superscript *n* for **x** and **z**, as we modeled trials as independent and identically distributed (i.i.d.). Further, we also drop the superscript m as the stimulus dependence will become apparent below.

To model temporal structure in the data, we put a first-order auto-regressive prior – also known as a linear dynamical system prior – on x_*t*_:

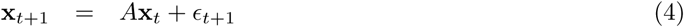

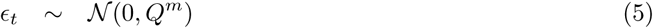

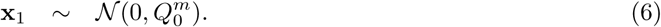

To allow for different levels of variability under the three experimental conditions *m* = 1, 2, 3 (corresponding to song, ML noise, and no stimulus, respectively), we introduce different covariance matrices *Q^m^* for the zero-mean innovations *ϵ_t_*. The factors in the first time step x_1_ have zero mean and covariance *Q_0_^m^*. We denote the PLDS model parameters as *θ*_PLDS_ = (*A*, *Q*, *C*) with *Q* = (*Q^1^, Q^2^, Q^3^, Q_0_^1^, Q_0_^2^, Q_0_^3^*).

Empirically, we found that the magnitude of the gain modulation **g** is small compared to the variability **z**. However, fitting PLDS models without the slow gain g to the same data resulted in the variability z to model slow non-stationarities in the data with time constants ≥ 1s. Therefore we included the **g**-term in the model to prevent these slow contributions from leaking into the fast variability **z**.

#### 2.4.3 Inference in the PLDS model

Given data **y** and the model parameters *θ*: = (*θ*_PSTH_, *θ*_PLDS_) we are interested in inferring the latent variables **x**. For the PLDS model with transfer function *f* (*x*): = log(1 + exp(*x*)), exact inference is intractable, and we therefore utilize Laplace inference, a general and widely used approximate inference method. For Laplace inference we determine the maximum a posteriori (MAP) value x:

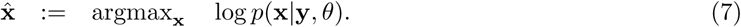

We optimize eqn. (7) by Newton’s method, which can be carried out efficiently in the PLDS model as the Hessian of the objective function is block tri-diagonal, and matrix equations involving the Hessian can therefore be inverted with computational time scaling linearly in *T*. The Laplace approximation *q*(x) ≈ *p*(x|y, *θ*) is then defined as a normal distribution *𝒩*(x|x̂, −H(x̂)^−1^) centered at x̂ with a covariance matrix given by the negative inverse Hessian −H(x̂)^−1^ of *logp*(x|y, *θ*) evaluated at x̂. Further details on Laplace inference in dynamical system models are given e.g. by Paninski et al. (2010).

#### 2.4.4 Model selection on number of latent factors

For each data set, we chose the dimension *d* of the PLDS model in the following way. We fitted model parameters *θ* for each latent dimension *d* = 1,…, 10 as described below and picked the dimension *d* which maximized the five-fold cross-validated log-likelihood log *p*(**y**_test_) on held-out test data under the model. Calculating this quantity for the PLDS model is intractable, so we replaced the marginal log-likelihood by the following “variational” lower bound which can be evaluated more easily (dropping the subscript of **y**_test_ for simplicity):

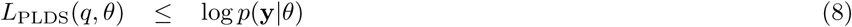

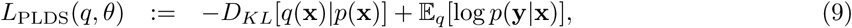

where 𝔼_*q*_ [∙] denotes the expected value under a distribution *q*. This bound holds for any distribution *q* (Emtiyaz Khan et al., 2013), but we seek to evaluate it using the Laplace approximation for *q*. The first term of eqn. (9) is given by the Kullback-Leibler divergence between *q*(**x**) and the linear dynamical system prior *p*(**x**). As both distributions are normal, one can evaluate this term analytically as a function of the model parameters *θ* as well as the mean and covariance matrix of *q*. The second term of eqn. (9) is the expected value of the log-likelihood under the Laplace approximation. For our choice of the transfer function *f*, we cannot evaluate this term analytically. We therefore resort to numerical integration using the fact that the likelihood decomposes into a sum over stimuli, trials, neurons and time steps:

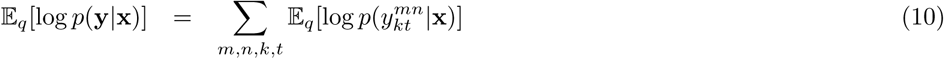

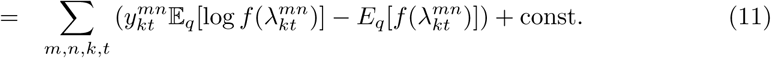

Under the approximate posterior *q*, the *λ_kt_^mn^* are normally distributed. Therefore, it is sufficient to evaluate the Gaussian integrals 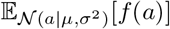 and 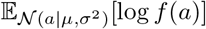 for different means *μ* and variances *σ*^2^. We tabulated these two functions on a two-dimensional grid over (*μ*, *σ*^2^) and approximated the actual values by linear interpolation.

#### 2.4.5 Estimation of PLDS model parameters

Given the collected spike count data **y**, we estimated the parameters *θ* = (*θ*_PSTH_, *θ*_PLDS_) in the following way. For simplicity, we first estimated the PSTH model parameters *θ*_PSTH_ as described in section 2.4.1. Then, keeping *θ*_PSTH_ fixed, we estimated the PLDS parameters *θ*_PLDS_, using the following slightly modified model. We add a stimulus dependent bias parameter *b_k_^m^* to eqn. (2):

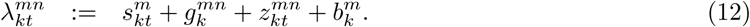

The rationale behind these additional parameters is the following. By construction, for different stimulus types (song vs. ML noise vs. silence) the Gaussian variables *z_kt_^mn^* have different variances. Due to the non-linear transfer function *f*, changes in variance of *z_kt_^mn^* will also cause changes in the mean of *y_kt_^mn^*, and not only its variability. Hence, we add biases *b_k_^m^* to compensate for such changes in mean of *y_kt_^mn^* caused by variance of *z_kt_^mn^*. After estimation of the model parameters *θ*_PLDS_ and **b** = (*b_k_^m^*)_m,k_, we eliminate the latter by absorbing it into the signal component **s**, i.e. *s_kt_^mn^* ← *s_kt_^mn^* + *b_k_^m^*. In general, this two-stage procedure might lead to sub-optimal estimates compared to jointly estimating *θ*_PSTH_ and *θ*_PLDS_. However, given its simplicity and the good model fits, this approach is well justified.

We estimated *θ*_PLDS_ and **b** using a variant of the expectation-maximization algorithm (Dempster et al., 1977), an iterative algorithm consting of an expectation (E-step) and a maximization (M-step) (Smith and Brown, 2003, Kulkarni and Paninski, 2007). In the E-step, we need to infer moments of the posterior distribution *p*(**x**|**y**) over the latent factors **x**; as the exact computation of these moments is intractable for the PLDS, we used the Laplace approximation described in section 2.4.3. The M-step for the PLDS model decomposes into two separate problems: first, updating the parameters *A, Q, Q*_0_ of the linear dynamical system prior, and second, estimating the loading *C* and biases **b** of the observation model eqn. (3). The former updates are essentially the same as described e.g. in Macke et al. (2011), Buesing et al. (2012), only slightly generalized to account for the different covariance matrices for different stimulus conditions. For updating the observation parameters, we need to find the new loading *C* and biases b that maximize the lower bound *L*_PLDS_ defined in eqn. (8) given the Laplace posterior approximation *q*(**x**) on the training data:

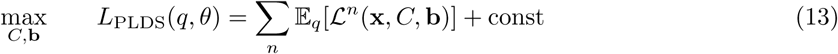

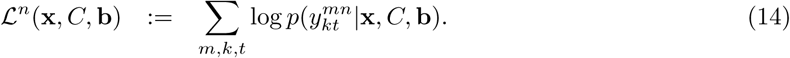

To circumvent the intractability of this problem, we used stochastic gradient ascent on *L* to update *C*, **b** (Bottou, 2010). We detail this approach for the loading matrix *C*; updates for **b** were analogous. Instead of exactly evaluating the cost function *L* and its gradient ∇_c_ *L* at each optimization iteration, stochastic gradients use a cheap stochastic approximation to these quantities, and perform well in practice. Let *C^l^* (and analogously *b_k_^lm^*) denote the estimate of *C* at iteration *l* of the inner loop of the M-step. We update the estimate in the following way:

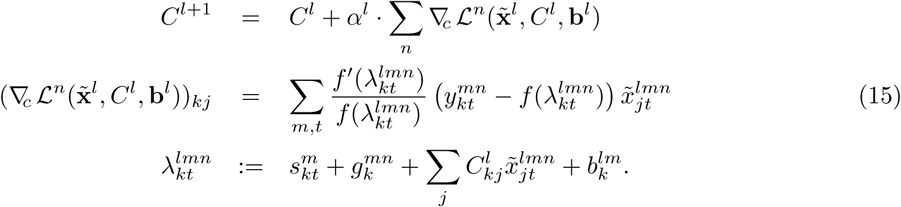

Here, x˜^*l*^ is a sample from the Laplace approximation to the posterior x˜^*l*^ ~ *q*(∙|y), with component *x˜_jt_^lmn^* for stimulus m on trial *n* in time bin *t* for factor *j*. As can be seen from eqn. (15), it is straightforward to evaluate the gradient ∇_*c*_ *𝓛^n^*, in contrast to *∇_c_ L.* The gradient with respect to *b_k_^m^* reads:

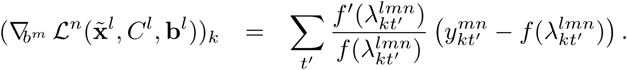

The decreasing learning rate α^*l*^ is given by:

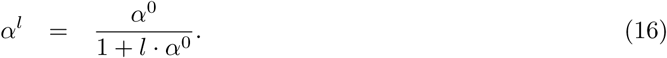

In each M-step we did 100 passes through the data set, i.e. *l* = 1,…, 100. We set *α*^0^ to depend on the iteration number *a* of the outer EM-loop according to *α*^0^ = (10^2^ + a)^−1.5^. Parameters were initialized using Exponential Family PCA (Collins et al., 2002).

#### 2.4.6 Decomposition of variability

Here we outline how we quantified the magnitude of the three model contributions to the hidden rate λ as in eqn. (2). We denote with 𝔼_*n*_[∙] the empirical average 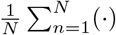 over trials *n* = 1,…, *N* and analogously 𝔼_*t*_, 𝔼_*m*_, 𝔼_*k*_; let 𝔼_*nm*_[∙] = 𝔼_*n*_[𝔼_m_[∙]], and 𝔼 denote the expectation over **x** and **z**. Using this notation, we can write the total covariance between λ of neuron *k* and neuron *j* averaged over stimuli, trials and time as:

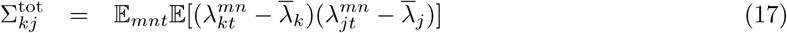

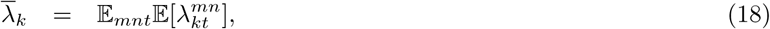

from which the following identity can be derived:

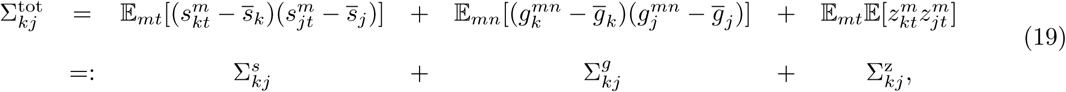

where 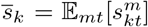 (and analogously g¯_*k*_). According to eqn. (19), the variability of the pre-intensities can be decomposed into a stimulus contribution Σ^*s*^, a contribution Σ^*g*^ due to the slow gain modulation and the shared noise contribution Σ^z^ modeled by the PLDS. Σ^*s*^ and Σ^*g*^ can directly be computed from eqn. (19) using the model parameters *θ*. The PLDS contribution can be computed from the parameters *θ*_PLDS_ in the following way:

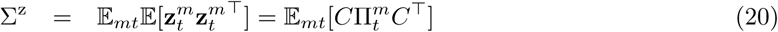

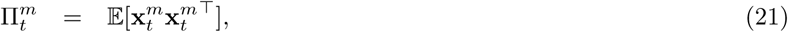

where Π*_t_^m^* is the prior covariance of the factors x_t_^m^ during presentation of stimulus m at time step *t*. The latter can be calculated using the standard recursion:

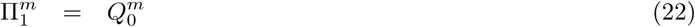

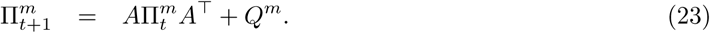

We define the magnitude of the stimulus drive for a neuron *k* as Σ*_kk_^s^*, and analogously Σ*_kk_^z^* and Σ*_kk_^g^*. We quantified the overall magnitude of the stimulus drive as the trace norm of Σ^*s*^ (scaled by the number of neurons *K*), i.e. *K*^−1^ trace[Σ*^s^*] = 𝔼_*mt*_[Σ*_kk_^s^*] (analogously for magnitudes of gain and shared variability). We also computed these magnitudes for the three experimental conditions individually by not taking the average 𝔼_*m*_.

#### 2.4.7 Coordinate transformation of PLDS model

It is known that many latent factor and state-space models (such as the PLDS) are non-identifiable, i.e. different model parameters can generate the same distribution over observed variables. For example, we can arbitrarily scale up the latent factors **x** by a scalar *α* if we scale down the loading matrix *C* by *α^−^*^1^ at the same time, leaving *p*(**y**|θ_PLDS_) unchanged. Buesing et al. (2012)shows that an arbitrary invertible linear transformation can be chosen to transform the latent factors, effectively choosing different coordinate systems to represent the latent factors. Here we make the following choice for the latent coordinate system to resolve this non-identifiability. We transform x such that the loading matrix *C* is orthogonal (*C*^T^*C* = *I*) and the averaged prior covariance matrix E*_mt_*[Π*_t_^m^*] is diagonal with decreasing entries on the diagonal. Furthermore, without loss of any generality we invert the signs of each latent factor *j* such that its corresponding column *C_:j_* of the loading matrix has at least as many positive entries as negative ones. This convention guarantees that different factors project to orthogonal subspaces (as *C*^T^*C* = *I*) and that factors are sorted with respect to their magnitude, i.e. the amount of variance of λ they capture. Using this convention, the latent factors are uniquely defined.

#### 2.4.8 Sign of loading matrices

For each PLDS model, we report the fraction of neurons which have a positive loading coefficient *C_k_*_1_ onto the first latent factor *x*_1*t*_. By definition, this factor captures the largest fraction of shared variability of the pre-intensity λ, and therefore neurons that have loading coefficients *C_k_*_1_ with the same sign are highly likely to have a positive noise correlation. We do a similar analysis for the signal and slow gain contributions Σ^*v*^, *v* ∈ {*s*, *g*}. To this end, we compute the first principal component *PC^v^* ∈ ℝ^*K*^ of Σ^*v*^ and report the largest fraction of neurons that have the same sign onto PC^*v*^.

#### 2.4.9 Quantifying dissimilarity between model contributions

We quantify the dissimilarity between two variance contributions *v*_1_ and *v*_2_ for *v*_1_, *v*_2_ ∈ {*s*, *z*, *g*} using the angle *ρ*(*v_1_, v*_2_) between their corresponding covariance matrices:

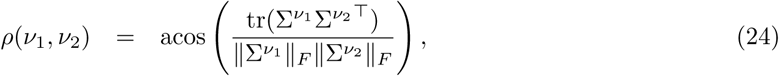

where || ∙ ||_F_ is the Frobenius norm. We compare *ρ* to the null distribution of angles between randomly oriented covariance matrices with the same eigenvalue spectra as Σ^*v*_1_^ and Σ^*v*_2_^. We can sample from this null by doing an eigenvalue decomposition of Σ^*v*_1_^ = *USU*^⊤^ and generating a surrogate covariance matrix 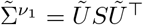, where *U˜* is a random matrix sampled from the uniform distribution over orthogonal matrices and computing the resulting angle *ρ*˜ (*v*_1_, *v*_2_) between Σ˜^*v*_1_^ and Σ^*v*_2_^.

#### 2.4.10 Autocorrelation functions and time constants

To quantify temporal continuity in the variance contributions, we computed their temporal autocorrelation functions *R*. For the signal component, we first computed for each neuron k the time-lagged covariance *R˜_k_^s^*:

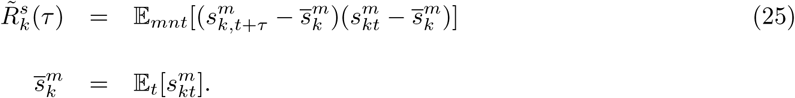

Normalizing *R˜_k_^s^*(*τ*) by *R˜_k_*(0)^−1^ yields *R_k_^s^*(*τ*). We report *R^s^*(*τ*) averaged over all neurons from all data sets. For the PLDS model, we computed the autocorrelation on the posterior distribution over the variability term **z**. We first computed the posterior mean 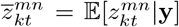. Based on this we compute *R^z^*(*τ*) in analogy to eqn. (25). This definition of *R^z^* also ignores the contribution of the posterior uncertainty. We found empirically that this contribution is small compared to the contribution of the posterior mean. We also qualitatively compared the posterior to the prior autocorrelation functions, finding only small differences.

We also characterize the temporal properties of the data by computing the time constant of the dynamical system prior that was learned when fitting the PLDS model parameters *θ*. The time constants *τ_i_* are related to the eigenvalues *e_i_* of the dynamics matrix *A* as:

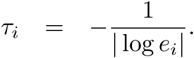

#### 2.4.11 Choice of transfer function

In most previous studies which applied latent variable models with Poisson observations to neural recordings, the transfer function was assumed to be *f* (*x*) = exp(*x*), as this choice offers multiple algorithmic advantages (Macke et al., 2011, Vidne et al., 2012, Paninski et al., 2010, Buesing et al., 2014). From a scientific viewpoint, it has been argued that neural response variability, especially in the visual system, can be modeled as a multiplicative gain modulation (Goris et al., 2014). Such a multiplicative noise model is equivalent to an additive noise source in the log-domain, and hence multiplicative gain modulation naturally falls within the class of PLDS models with exponential transfer function. For the data considered here, we found that the PLDS model with exponential transfer function could model the activity during stimulus presentation accurately. However, the model failed to capture the statistics of spontaneous activity: the variances and cross-covariances of the neural activities were substantially over-estimated (data not shown). We found that a PLDS with the transfer function *f* (x) *=* log(1 + exp(*x*)) did not have the same shortcoming and that it also improved the test log-likelihood significantly over a model with *f* (*x*) = exp(*x*) when trained exclusively on spontaneous activity. (Recently-developed methods (Gao et al., 2016) permit the nonlinearity to be estimated directly from data on a neuron-by-neuron basis; we leave this extension for future work.)

## 3 Results

### 3.1 Latent factor model captures shared variability

We fitted the PLDS model separately to each of the ten recording sessions from the deep region of the songbird A1. One example fit for a data set with 29 simultaneously recorded neurons is shown in Figure 1B. A first observation that can be made from this example is that, in contrast to the stimulus drive s_t_, the shared variability **z**_*t*_ has a simple structure. Namely, **z**_*t*_ is highly coordinated across neurons, to the effect that the firing rate fluctuations of most neurons increase (or decrease) at the same time. This is a result of the fitted models being “low-rank”, i.e. a large fraction of the variance of **z**_*t*_ is captured by the first latent factor *x*_1,*t*_ (analogous to the first principal component in a PCA analysis). This can also be seen from Figure 1C: the time series of the dominant first factor *x*_1,*t*_ is highly similar to that of the summed population noise; we defined the latter as the total number of spikes summed across the recorded population minus the total number of spikes predicted by the PSTH (Macke et al., 2011, Okun et al., 2015). Furthermore, *x*_1,*t*_ summarizes the trial-to-trial variability across the population for repeated presentations of the same stimulus (see Figure 1D).

The low-rank nature of the shared variability **z**_*t*_ in the example data set of Figure 1B was found in all of the ten datasets we analyzed (Figure 2A). Although the optimal number of latent factors (chosen to maximize the likelihood evaluated on held-out test data) ranged from 1 to 7 for different data sets, at least 88% of the variance in **z**_*t*_ is captured by the first latent factor *x*_1,*t*_ in all data sets. Averaged over all data sets, the first component captures more than 96% of the shared variability of the full model. This means that shared variability in our recordings is dominated by a single, unobserved noise contribution, and not e.g. by independent noise terms for pairs of neurons. In contrast to **z**_*t*_, the stimulus drive **s**_*t*_ is not low-rank: its variance is distributed across many dimensions (Figure 2A), resulting in the first principal component capturing on average only 31%, and at most 53% across all data sets. We also found that the parsimonious description of the the population variability provided by the model crucially depended on the nonlinearity of the transfer function *f*: linear factor models failed to discover the low-rank structure of the data. We show the results of directly applying PCA to the raw population spike count variability, which we computed by subtracting the population PSTH from the spike counts **y**_*t*_ (Figure 2A).

**Figure 2:**
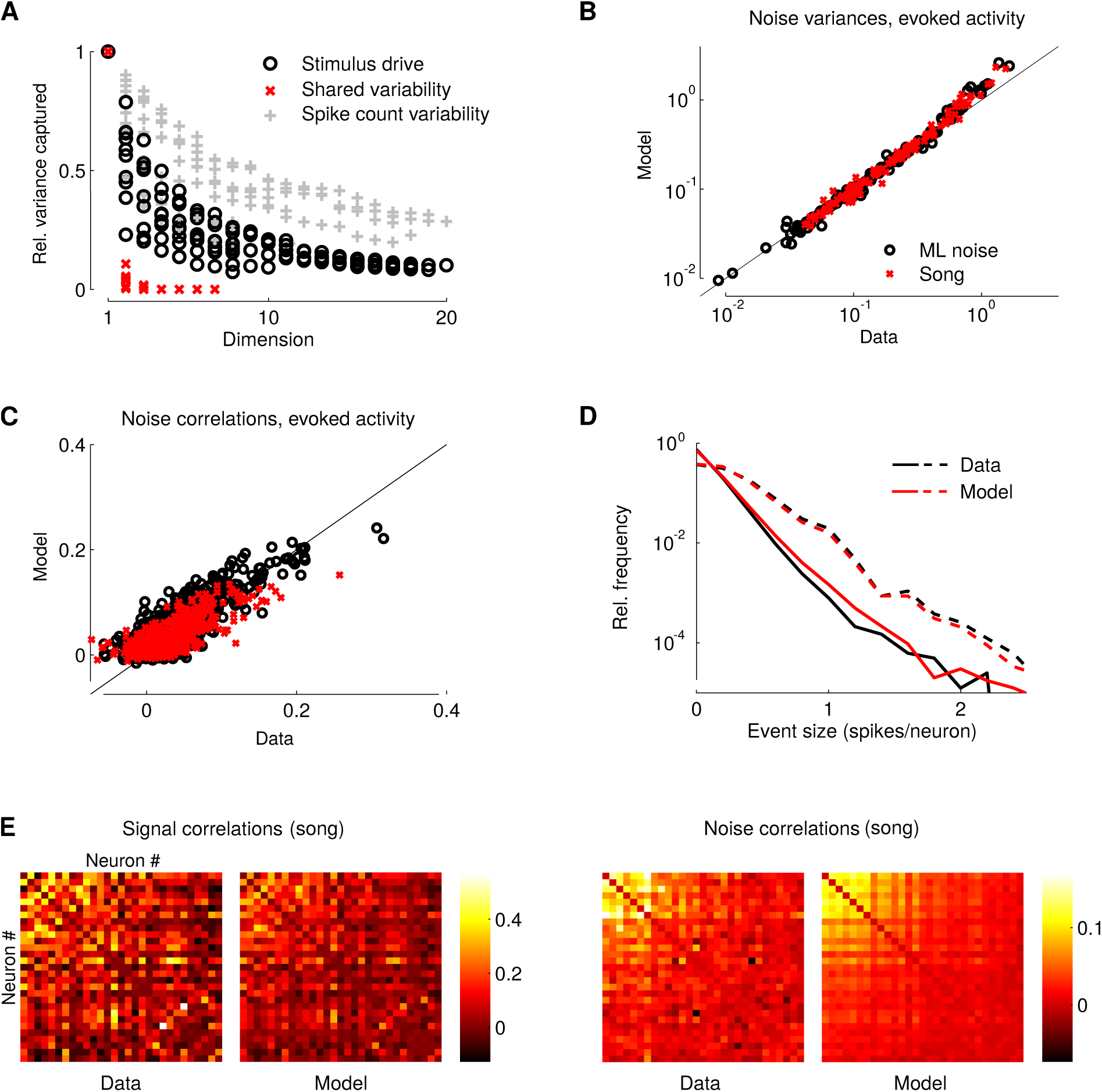
Low-rank latent factor model faithfully captures songbird A1 population recordings. (**A**) Eigenvalues of the covariance matrices of the stimulus drive **s**_*t*_ (black) and the population variability **z**_*t*_ (red) of the PLDS model ordered with respect to their magnitude, pooled over all data sets from deep region L3 (scaled such that largest eigenvalue for each data set is 1). This shows that variance of the signal drive **s**_*t*_ is distributed over many dimensions, whereas the shared variability **z**_*t*_ is low-rank. PCA on the raw population spike count variability (**y**_*t*_ minus population PSTH) fails to uncover the low-dimensional structure in population variability. (**B**) Noise variances of all recorded cells from deep region L3 (double logarithmic plot) computed from raw data (horizontal axis) and PLDS model fits (vertical axis). Each marker represents a neuron under one corresponding stimulus condition (ML noise: black, song: red). Black line indicates the diagonal. (**C**) Same as (**B**) but for all pairwise noise correlations. Each marker represents a pair of simultaneously recorded neurons under the corresponding stimulus condition. (**D**) Empirical distributions of summed population spike count (number of spikes in 50 ms windows averaged across population) computed from raw data (black dashed) are matched by that from surrogate data sampled from the PLDS model (red dashed). Solid lines are distributions of the population spike count after subtracting the PSTH for each cell; this shows that the PLDS model faithfully captures the statistics of the summed population response variability. (**E**) Example signal and noise correlation matrices computed from raw data and PLDS surrogate data for the same recording session as shown in Figure 1.

In spite of the simple structure of the fitted PLDS models, they capture the recorded data well. We compared various summary statistics computed from the raw data with those computed from surrogate data generated by the models. We find that the sampled surrogate data reproduces average single cell properties (mean firing rate and PSTH, data not shown). The PLDS model also captures the variability in evoked responses faithfully (Figure 2B-E), such as single cell noise variances (Figure 2B). Furthermore, the model explains 81% of all pairwise noise correlations computed from both stimulus conditions together, while also matching noise correlations under the two different stimulus conditions separately (Figure 2C). Figure 2E shows the noise correlation matrix under the song condition for an example data set together with the corresponding signal correlation matrix, illustrating the match between experimental and model surrogate data. Beyond single cell and pairwise noise statistics, the model also captures the statistics of the overall population activity; for example, Figure 2D shows that the model matches the observed empirical distribution over the summed population spike count defined above, also capturing rare events with relative frequency of 10^−4^ (which occur at most 2-3 times in the recorded data). On held-out test data, we find that averaged across all data sets, the PLDS model provides 4.5 ∙ 10^−2^ more bits of information per spike compared to a model without the variability term **z**_*t*_. This performance is comparable to the highest values previously reported for population variability models applied to data from primate motor cortex (Pachitariu et al., 2013, Gao et al., 2015). Taken together these findings establish that a large fraction of the population shared variability in the deep region of the songbird A1 is explained by a single latent factor.

### 3.2 Shared variability is stimulus dependent in the deep region of avian A1

Having established that the PLDS model provides a faithful statistical description of the recorded data, we quantified and characterized the three contributions to the population response in the model, i.e. the signal drive **s**_t_, gain modulation **g** and variability **z**_*t*_. First, we used the model to investigate population aspects of song selectivity, by comparing the magnitude of the model contributions between the song and ML noise stimulus conditions. We quantified the magnitudes of the individual terms by the trace norm of their respective covariance matrices normalized by population size *K*, e.g. for the stimulus drive by tr[Cov(s)]/*K*; this can be conveniently interpreted as the average power of the signal drive per neuron. We found that in the deep region of avian A1 the stimulus drive is larger for song than for ML noise stimuli (Figure 3A). In addition to the signal drive, the model captures a considerable contribution of shared variability in the data, which for ML noise is on the same order of magnitude as the stimulus drive, indicating strong, shared fluctuations in this neural population. Notably, the population variability is substantially larger (by roughly a factor of 2) under the ML noise compared to the song stimuli, showing that the stimulus class has a large influence on the shared variability of local populations of simultaneously recorded neurons. Finally, we find that the slow gain modulation g only captures a small amount of variance in the data and is of roughly the same magnitude under song and ML noise stimuli. In Figure 3A the lighter shaded areas of the bars represent the fractions of variance of each term that are captured by a one dimensional approximation, i.e. the first principal component or the first factor *x*_1*,t*_ respectively. For the signal drive **s**_*t*_ as well as the gain modulation **g**, less than half of the variance is captured by the first component, indicating that these two effects are heterogeneous, with a much larger effective dimensionality than the shared variability **z**_*t*_ (c.f. Figure 2A).

**Figure 3:**
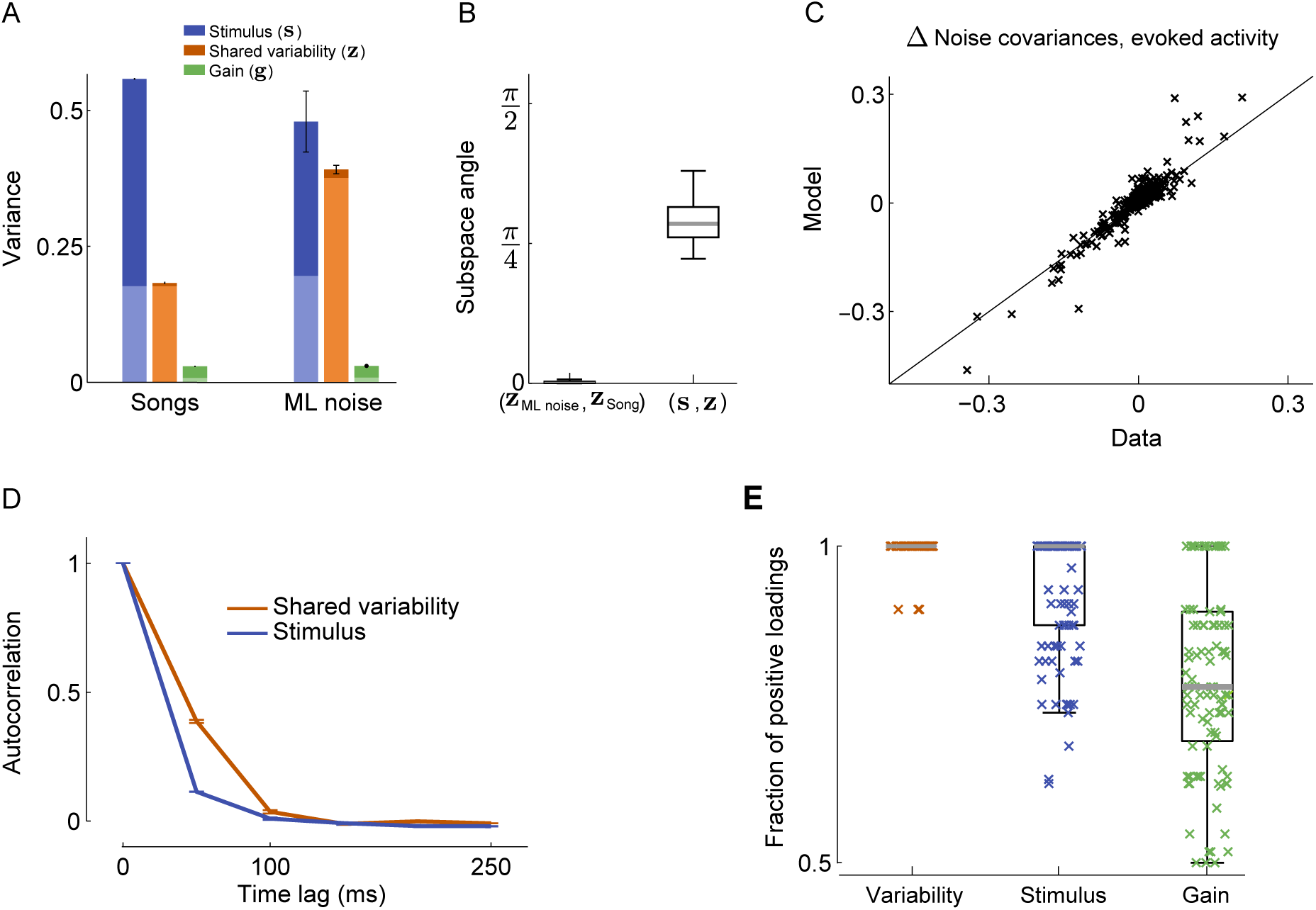
Shared variability is stimulus dependent in the deep region of avian A1. (**A**) Magnitude of stimulus drive **s**_*t*_, population variability **z**_*t*_ and gain modulation g for both stimulus conditions quantified by the trace norm of their respective covariance matrices and normalized by the number of neurons. While the stimulus drive is larger for conspecific song stimuli, the corresponding population variability is substantially reduced compared to ML noise stimuli. Lower part (light region) of each bar indicates share of variance that is captured by a one-dimensional approximation, highlighting that the shared variability but neither stimulus drive nor gain modulation are low-rank. (B) Angle between population variability **z**_*t*_ under song and ML noise conditions is close to zero, while angle between population variability **z**_*t*_ and stimulus drive **s**_*t*_ within a condition is substantially larger. (**C**) Difference of the noise covariances (including variances) between song and ML noise conditions for all simultaneously recorded pairs of neurons from L3; computed from data (horizontal axis) and from PLDS model fit (vertical axis). Black line indicates diagonal. (**D**) Autocorrelation functions of signal drive **s**_*t*_ and shared variability **z**_*t*_ averaged over all data sets. Error bars for (**A**) and (**E**) are given by standard deviation of mean over five-fold cross-validation. (**E**) Fraction of cells that have the same sign of loading coefficient onto a onedimensional approximation to the terms **s**_*t*_, **g**_*t*_ and **z**_*t*_ respectively. Each cross represents one model fit on one data set. This illustrates that the population variability **z**_*t*_ increases (and reduces) the firing of almost all cells together, whereas this is less the case for **s**_*t*_ and **g**_*t*_.

In addition to the magnitude and effective dimensionality of the terms **s**_*t*_, **g** and **z**_*t*_, we analyzed their similarity by comparing the directions they spanned in the firing rate space (or more precisely in the *K*-dimensional pre-intensity space). We quantified this by computing the subspace angle, which lies between 0 (perfect alignment) and *π*/2 (orthogonality, for details see Materials and Methods). Figure 3B shows that the angle between the dominant subspace of the shared variability term **z**_*t*_ under song stimuli and under ML noise stimuli is small: for all data sets the angle was less than 0.1, indicating that the shared variability **z**_*t*_ is highly aligned across stimulus classes when compared to the null hypothesis of random orientation with the same spectra (*p <* 10^−10^,Wilcoxon rank sum test). In comparison, the median angle between shared variability **z**_*t*_ and stimulus drive **s**_*t*_ was 0.90. (Although substantially larger than that for **z**_*t*_ across stimulus classes, the contributions **z**_*t*_ and **s**_*t*_ to the population response are still significantly more aligned than expected under the null hypothesis; p < 10^−10^,Wilcoxon rank sum test.) Hence, shared variability is much more similar across stimulus conditions than it is to stimulus drive within the same condition. This implies that the stimulus class largely changes the magnitude of shared variability and not its direction in the *K*-dimensional pre-intensity space. We verified that this surprisingly simple interpretation derived from the model is indeed an accurate description of the raw data. Figure 3C shows the difference of all elements of the noise covariance matrices under song and ML noise stimuli. The PLDS model prediction, consisting of a simple change of magnitude of shared variability and not its direction, matches the data to a large extent (R^2^ = 0.87).

We next analyzed the temporal properties of the essentially one-dimensional shared variability term **z**_*t*_, by computing the autocorrelation of **z**_*k,t*_ for each neuron *k* and averaging over the population. Figure 3D shows that the autocorrelation of the shared variability decays after roughly 200 ms. This is consistent with the time constants of the dynamical system prior of the latent factors **x**_*t*_ learned by fitting the PLDS to the data: the average time constant (pooled across data sets) is 105± 114ms (mean ± std. deviation). Taken together with the above finding that the gain modulation **g** is small, we concluded that most network-level variability in these recordings takes place on fast time scales below 200 ms.

Although largely temporally unstructured, the shared variability term **z**_*t*_ seems to be highly structured with respect to its influence on the recorded neurons (see Figure 1B). Indeed, we found that almost all neurons *k* have a positive loading coefficient *C_k_*_1_ onto the first factor *x*_1,*t*_ of **z**_*t*_ (see Figure 3E). This means that the network activity fluctuations captured by *x*_1,*t*_ cause a temporally coherent activation and deactivation of the majority of recorded neurons. This finding is also in accordance with the fact that most noise correlations in the data are positive (see Figure 2C). For comparison, we also analyzed the signs of loadings onto the first principal components of the stimulus drive **s**_*t*_ and gain modulation **g**. We found a similar if less pronounced effect for the stimulus drive, indicating that **s**_*t*_ also induces mostly positive signal correlation, whereas the gain modulation influence is largely unstructured.

Taken together, the PLDS model-based analysis presents the following parsimonious description of the shared population variability: the stimulus identity modulates the magnitude of variability of a very simple (i.e., approximately one-dimensional) network state, modeled by the first latent factor. The activity of most neurons are simultaneously increased and decreased, reflecting the fact that observed noise-correlations are positive to large extent.

### 3.3 Latent shared variability is spatially structured in the deep region of songbird A1

As discussed above, we found a strong stimulus effect on the level of shared variability in the deep region (Figure 3), whereas the same analysis discovered little shared variability in the intermediate regions for either stimulus class (data not shown). Going beyond this spatially coarse description by region, we were interested in the detailed spatial organization of the latent variability in the deep region L3, in particular close to the anatomical boundary between the intermediate and deep regions.

We histologically determined the approximate location of the polytrode recording sites, yielding an estimate of the spatial location for each recorded neuron (see Figure 4A). A first hint at a corresponding, fine-grained spatial structure of shared variability can be observed in Figure 1B (bottom panel) and Figure 2E. For these plots we sorted the neurons with respect to spatial location. This sorting uncovered a clear spatial gradient in the strength of shared variability in the data as well as in the corresponding model fit. To characterize this effect in greater detail, we quantified the strength of shared variability for each cell *k* by computing the variance Var(*z_k_*,_*t*_) that the variability term **z**_*t*_ captures in the firing of cell *k*. We call this quantity the magnitude of the shared variability in cell k. In Figure 4B we pooled neurons across all data sets and plotted the magnitude of shared variability as a function of recording location. A clear, monotonic increase in shared variability as a function of spatial location can be observed, and both stimulus conditions exhibit significantly increased shared variability at 0.5 mm distance from the intermediate regions (*p <* 3 ∙ 10^−4^, Wilcoxon rank sum test). Consistent with the results above, shared variability was larger for ML noise stimuli compared to song stimuli across the whole spatial range. However, the difference was significantly larger deep in L3 compared to the intermediate region (p < 2 ∙ 10^−4^, Wilcoxon rank sum test). Hence, we see a gradual increase in song selectivity, i.e. a relative increase of shared variability under ML noise compared to song stimuli. In contrast to these results, we did not find a clear spatial trend for the signal drives (Figure 4C), i.e. the median magnitude of s at 0.5 mm depth is not significantly different from that in the input region (*p* > 0.2, Wilcoxon rank sum test). For the gain **g**_*t*_ we observed a small but significant increase of the median magnitude from the intermediate region to deep L3 by about 30% (*p* < 0.02, Wilcoxon rank sum test; data not shown).

**Figure 4:**
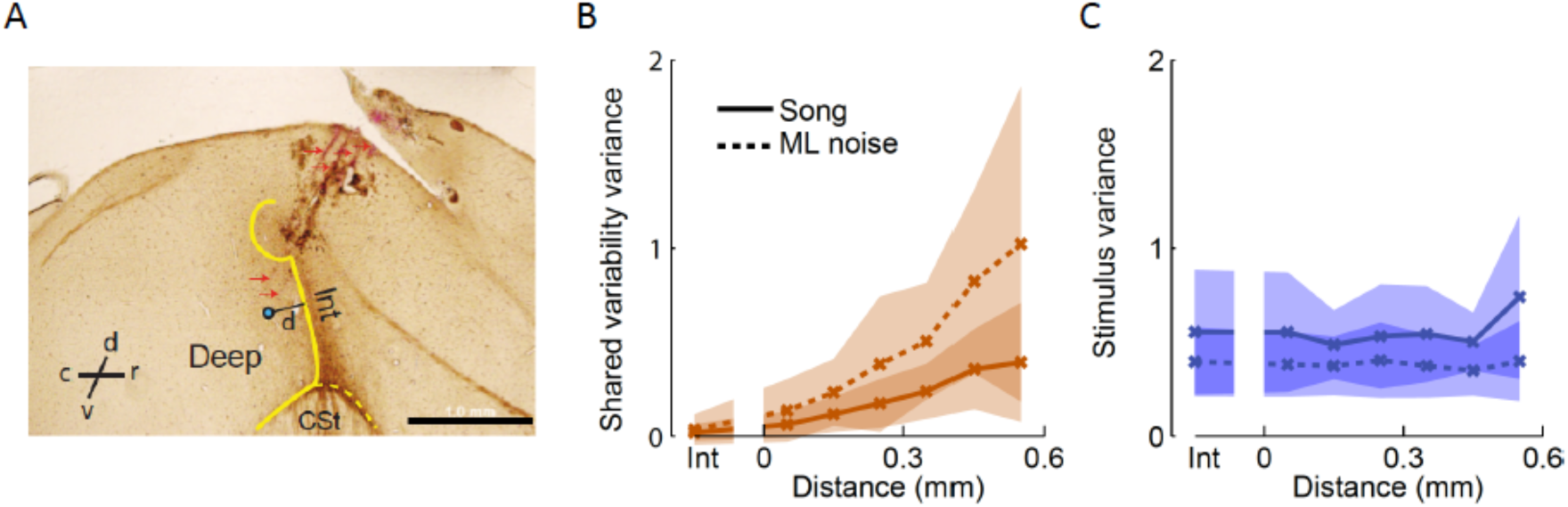
Shared variability is spatially structured in the deep region of songbird A1. (**A**) Parasagittal section of the avian auditory cortex. The border between the intermediate and deep regions is indicated in yellow and the blue dot indicates the location of an example recording site. The distance between a recording site and the intermediate region was defined as the minimum distance to the yellow contour. Purple arrows indicate electrode tracks (CSt: Caudal striatum). (**B**) Magnitude of shared variability increases with distance from the intermediate region, and the rate of increase is larger for ML noise stimuli. This implies increasing song selectivity in shared variability as a function of distance from the anatomical boundary. Each data point shows the average across units located at the same distance (binned at 0.1 mm) and error bars indicate standard deviation across cells. (**C**) Same as (**B**) but for magnitude of signal drive **s**_*t*_. In contrast to shared variability, a clear relationship with distance is not evident.

It is worth noting that the latent factors in the PLDS model allowed us to construct these spatial maps of shared variability. Instead of looking at pairwise correlations (which would yield pairs of locations), the PLDS allowed us to define a magnitude of shared variability per neuron, which can be thought of as a coupling strength of each neuron to an unobserved network state.

### 3.4 Stimulus onset quenches shared variability

Having established that population response variability of evoked activity is spatially structured and has an essentially one-dimensional form, we next sought to characterize its relation to spontaneous activity. To this end, we focused on the recorded neural activity in time windows of 500 ms preceding the onset of each stimulus presentation; these epochs of spontaneous activity were included in the data sets to which the PLDS models were fitted. We found that the PLDS model could account for the statistics of spontaneous activity reasonably well: the model was able to capture R^2^ = 73% of the noise correlations during spontaneous activity, although a tendency to slightly over-estimate their magnitude is visible (see Figure 5A). We found that spontaneous activity has a simple statistical structure: As the signal drive term is zero by definition, spontaneous activity is exclusively captured by **z**_*t*_ (and gain modulation g), which again was found to be essentially one-dimensional (see Figure 5C). Furthermore, in the pre-intensity domain spontaneous activity is highly aligned to the shared variability during stimulus presentation (Figure 5D). Previous theoretical as well as experimental studies have argued that the distribution of spontaneous activity patterns should equal the distribution of activity patterns evoked by sensory stimuli, when averaging the latter over stimulus ensembles occurring under natural conditions (Ackley et al., 1985, Berkes et al., 2011, Buesing et al., 2011). However, we found spontaneous activity to be substantially different from evoked activity. Evoked activity induces activity patterns with high effective dimensionality (Figure 2A), whereas spontaneous activity in our data was approximately one-dimensional (Figure 5C). Furthermore, the latter was much closer to shared variability during evoked activity in terms of subspace angle, as can be seen from Figure 5D. Finally, note that the stimulus ensemble used in this study was highly limited compared to the diversity of auditory stimuli to which the birds are generically exposed. The observed dissimilarity of spontaneous activity from evoked activity would presumably further increase if compared to evoked patterns under richer stimuli.

**Figure 5:**
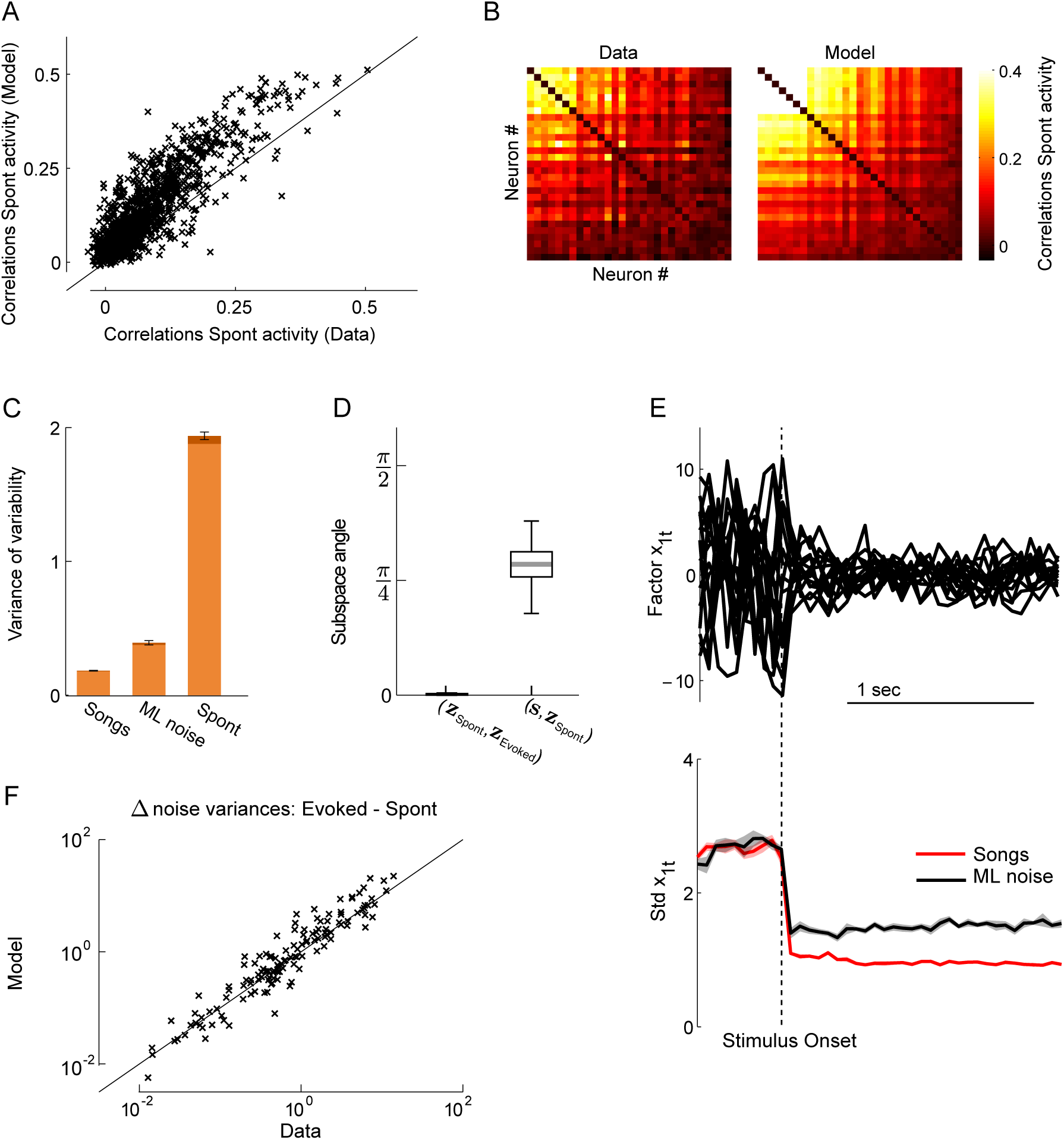
Shared variability of evoked activity is highly similar to spontaneous activity. (**A**) Noise correlations during spontaneous activity for all pairs of simultaneously recorded neurons from region L3, computed from raw data (horizontal axis) and PLDS model fits (vertical axis). Black line indicates diagonal. (**B**) Example noise correlation matrices during spontaneous activity computed from raw data and PLDS model fit for same data set as shown in Figure 1. (**C**) Magnitude of shared variability **z**_*t*_ during spontaneous activity. Lower, brighter part of the bar indicates variance captured by first latent factor *x*_1,*t*_ indicating that shared variability during spontaneous activity is low-rank. For comparison the magnitudes during ML noise and song stimuli are also shown (same data as in Figure 3A). (**D**) Left: Angle between shared variability z_t_ during spontaneous activity and during evoked activity. Right: Angle between shared variability **z**_*t*_ during spontaneous activity s_t_ and stimulus drive. (E) Top: Inferred first factor *x*_1,*t*_ (posterior mean) around stimulus onset for all 15 presentations for one song stimulus (same example data set as B). Bottom: Magnitude of first factor *x*_1,*t*_ (square root of posterior second moment) around stimulus onset, pooled over all song and ML noise stimuli for same example data set. (**F**) Differences in single cell noise variance between spontaneous and evoked activity (logarithmic scale) for all recorded neurons from L3, computed from data (horizontal axis) and PLDS model (vertical axis). Black line indicates diagonal.

The above analysis shows that spontaneous activity is highly similar to shared noise variability during stimulus presentation, but with larger magnitude. This finding is in agreement with previous findings that variability of single cell activity is in general substantially reduced by the onset of a stimulus or a task (Churchland et al., 2010). However, this effect has been mostly studied on the level of single cells, whereas the PLDS model allowed us to investigate the network structure as well as the temporal dynamics of the effect. We can visualize the shared network variability on single trials by plotting the first latent factor *x*_1_*,_t_* as a function of time. The results are shown in Figure 5E around the time of stimulus onset for all 15 presentations of one song stimulus in one recording session. It can be seen that the large variability of the latent factor is rapidly suppressed after stimulus onset, with the effect of “quenching variability” acting on a time scale of 50 ms or faster in this data set. This is also clearly visible when aggregating data across all stimuli (Figure 5F) from the same recording session. We observed similar results in all other data sets (not shown). Furthermore, we can show that the PLDS model correctly accounts for the changes of single cell noise variability following stimulus onset (R^2^ = 0.88, Figure 5G). Hence, using the PLDS model, quenching of single cell noise variability in this data can be explained to large extent as a result of reduced shared variability around stimulus onset.

## 4 Discussion

Using a latent factor model of population responses, we found that shared variability in populations of neurons in the songbird auditory cortex is fast (time-scale ≈ 100 ms) and that it modulates neural activity in a highly coordinated way, to the effect of synchronously increasing and decreasing the activity of most cells. Our main finding is that that a large fraction of this effect can be captured by a single degree-of-freedom latent factor, which can be interpreted as a shared mode of network activity, whose magnitude is strongly reduced by the onset of auditory stimulation, with greater reductions for conspecific song stimuli compared to artificial noise stimuli.

Cohen and Maunsell (2009) have shown that visual attention, widely believed to be a top-down phenomenon, reduces noise correlation in primate V4. Analogously, selective attention of birds to conspecific songs compared to artificial noise stimuli could be responsible for the observed reduction in shared variability; however, further experiments would be necessary to explore this hypothesis. It has recently been suggested that the reduction of pairwise correlations observed by Cohen and Maunsell (2009)can be accounted for by a low-dimensional latent factor model comparable to ours (Rabinowitz et al., 2015).

Using another approach similar in spirit to ours, Okun et al. (2015)studied the relation of single cell activity to the average activity of a population of simultaneously recorded, nearby neurons (the so-called population activity). They quantified this relation with a parameter for each cell termed population coupling (PC), which is essentially given by the cross-correlation coefficient between single cell and population activity. It was pointed out that PCs and loading coefficients of a Poisson factor model (Pfau et al., 2013) (similar to the one studied here) are conceptually related and show high degrees of numerical agreement. One major difference between our approach and that of Okun et al. (2015)is that we develop an explicit generative model of the unobserved influences which cause shared variability; this allows us to characterize their complexity (i.e. the number of latent factors), time scales, and magnitudes relative to the signal drive. Nonetheless, our finding that the correlation structure in this songbird auditory structure has an essentially one-dimensional structure is consistent with the finding in Okun et al. (2015) that a one-dimensional signal dominates the correlation structure in mammalian visual cortex (though see Lin et al. (2015)for an elaboration of this model).

More generally, there is rapidly increasing interest in investigating and characterizing latent structures from multi-cell data — e.g. estimating correlation patterns from partially observed populations or inferring common inputs from high-dimensional spike recordings — by applying sophisticated models similar to the PLDS model used here (Paninski et al., 2010, Macke et al., 2011, Buesing et al., 2014, Köster et al., 2014, Ecker et al., 2014, Gao et al., 2015, 2016), and we anticipate continued further development and applications of this modeling technology in the years to come.

## Acknowledgments

This work was supported by grants from the Simons Foundation (SCGB 325171 and 325233 to LP and JPC), grant ONR N00014-14-1-0243 (LP), and a Sloan Research Fellowship to JPC.

